# The effects of epigallocatechin gallate and caffeine on risky decision-making

**DOI:** 10.1101/2021.04.26.441489

**Authors:** A. E. Liley, H. Joyner, D. B. K. Gabriel, N. W. Simon

## Abstract

Epigallocatechin-3-gallate (EGCG) and caffeine are the two primary compounds found in green tea. While EGCG has anxiolytic and anti-inflammatory effects, its acute effects on cognition are not well understood. Furthermore, despite widespread green tea consumption, little is known about how EGCG and caffeine co-administration impact behavior. Here, we investigated the effects of multiple doses of either EGCG or caffeine on a rat model of risk-taking. This was assessed using the risky decision-making task (RDT), in which rats choose between a small, safe reward and a large reward with escalating risk of mild footshock. Rats were tested in RDT after acute systemic administration of EGCG, caffeine, or joint EGCG and caffeine. EGCG caused a dose dependent reduction in risk-taking without affecting reward discrimination or task engagement. Caffeine did not impact risk taking, but elevated locomotor activity and reduced task engagement at high doses. Finally, exposure to both EGCG and caffeine had no effect on risk-taking, suggesting that low-dose caffeine is sufficient to mask the risk-aversion caused by EGCG. These data suggest EGCG as a potential therapeutic treatment for psychological disorders that induce compulsive risky decision-making.

## Introduction

Green tea is a widely-consumed non-fermented tea derived from leaves of the *Camellia sinensis* tree (Cabrera et al., 2006; Singh et al., 2011). The polyphenols found in green tea have been recognized for their medicinal properties in the prevention of multiple diseases, including cancer, cardiovascular and neurological diseases, and diabetes (Khan and Mukhtar, 2019). In addition, green tea has been shown to have antibacterial, antiviral, and anti-inflammatory effects (Zhang et al., 2018), promote bone health (Shen et al., 2011), lower blood pressure (Negishi et al., 2004), and aid in weight loss (Kovacs et al., 2004).

The most abundant psychoactive ingredients in green tea are caffeine and a catechin called epigallocatechin-3-gallate (EGCG; Singh et al., 2011). EGCG is a natural phenol and flavonoid that suppresses inflammation, carries chemo-preventative effects, and may contribute to reduction of carcinogenesis (Cabrera et al., 2006; Liu et al., 2016; Kochman et al., 2020). Previous research has also shown that EGCG exerts modulatory effects on GABA_A_ receptors, likely at the benzodiazepine active site (Campbell et al., 2004; Vignes et al., 2006). This effect has been observed in mice with EGCG causing anxiolytic and amnesic effects comparable to the benzodiazepine chlordiazepoxide (Vignes et al., 2006). Human research has reported lower prevalence of depressive symptoms in elderly Japanese populations that could potentially be due to inhibition of the hypothalamic–pituitary–adrenal axis following higher consumption of EGCG found in green tea (Zhu et al., 2012).

Despite the prevalence of green tea consumption, the cognitive effects of EGCG are not well-established. EGCG has shown potential as a cognitive therapy in rodent models of neural disorders. In a rat model of brain damage caused by cerebral ischemia, EGCG reduces long-term memory loss by reducing oxidative stress and neuro-inflammation (Wu et al., 2012). EGCG has been shown to attenuate cognitive impairment in rodent models of Alzheimer’s Disease, improving spatial learning and long-term recognition memory in PS2Tg2576 mice (Kim et al., 2019), and enhancing working memory and spatial learning in APP/PS1 transgenic mice (Bao et al., 2020; Ettcheto et al., 2020). Furthermore, recognition memory and spatial working memory deficits are alleviated by green tea extract in genetic models of Down Syndrome (Feki and Hibaoui, 2018; De Toma et al., 2019).

The other major component involved with green tea is caffeine, which naturally occurs in tea leaves, coffee beans, and the cacao plant, functioning as an insecticide (Lee et al., 2009). As a stimulant, caffeine has reinforcing and psychostimulant effects similar to other stimulants (e.g., amphetamines, methylphenidate), inducing hyperactivity, improving mood, and increasing concentration (Diller et al., 2008). Caffeine acts as a non-selective antagonist at adenosine A_1_/A_2A_ receptors, which results in increased transmission of dopamine and norepinephrine in the nucleus accumbens (a region that plays a vital role in the dopaminergic reward system; Cardinal et al., 2001; Cauli and Morelli, 2005; Schmidt et al., 2001). Caffeine has also been shown to enhance cognitive performance and improve long-term memory with both single doses and repeated use in humans (Smit and Rogers, 2000; de Mejia and Ramirez-Mares, 2014). In a rat-model of decision-making, acute caffeine reduced delay discounting/impulsive choice, which is consistent with the effects of other stimulant drugs (Winstanley et al., 2006; Diller et al., 2008; Shiels et al., 2010; Rajala et al., 2015). Additionally, caffeine increases both physical and cognitive effort exertion during decision-making, although this enhancement was only observed in subjects with a preference for non-effortful options (Cocker et al., 2012; SanMiguel et al., 2018).

Despite the established health benefits of green tea, little is known about the effects of EGCG independently or with caffeine on decision-making. Risk-taking in the face of negative consequences is elevated in several psychiatric disorders (Orsini and Simon, 2020); therefore, it is critical to identify compounds that alleviate excessive risk-taking with low abuse potential that do not gross cognitive or motivational impairment. Risk-taking can be assessed in a rat model using the Risky Decision-making Task (RDT), which measures preference between a small, safe reward and a larger, risky reward accompanied by a chance of punishment (mild foot shock) which escalates throughout the session (Simon et al., 2009). This models situations in which decisions of greater economic value are sometimes accompanied by the risk of consequences (Orsini et al., 2019). Here, we tested the acute effects of systemic EGCG or caffeine on risky decision-making in rats. Then, we evaluated the effects of jointly administered EGCG and caffeine on risky decision-making.

## Materials and Methods

### Subjects

Male Long-Evans rats (n = 10, Envigo) were obtained at approximately 200 days post-natal. Subjects were individually housed and kept on a reverse 12-hour light/dark cycle with lights off at 7:30 am. Animals were food restricted to 90% of free feeding baseline weight to increase motivation and reward pursuit in behavioral tasks. All protocols were approved by the University of Memphis Animal Care and Use Committee.

### Behavioral Apparatus

Behavior and decision-making processes were measured in MedAssociates (FairFax, VA) operant conditioning chambers housed in soundproof cubicles. All chambers were equipped with one retractable lever on either side of an illuminable food trough with recessed photobeam for entry tracking, a pellet dispenser, metal shock grate, and photobeam sensors for locomotion tracking. Sugar pellets were obtained from Bio-Serv (Flemington, NJ) and used as a behavioral reinforcer.

### Instrumental Shaping

Shaping procedures to enable acquisition of RDT were similar to previous reports (Gabriel et al., 2018; Orsini and Simon, 2020). The day before training began, rats were given a handful of sucrose pellets (Bioserv, NJ) in their home cage to reduce neophobia. Day one consisted of Magazine training, during which 1 pellet was delivered into the trough every 30 ± 10 s for 38 trials to establish an association between the trough and pellet delivery. On the next day, subjects were trained to press a single lever for pellet delivery, followed by training on the other lever in subsequent days until 50 reinforcer pellets were obtained from each. Levers were presented in counterbalanced order across subjects. Then, rats were trained to nose poke into an illuminated food trough to trigger extension of a single lever in pseudorandom order (the same lever was never presented more than twice in a row). Each lever press resulted in delivery of one pellet and retraction of the lever, followed by a 10 ± 5 intertrial interval. Sessions were repeated until successful completion of 35 presses on each lever.

### Risky Decision-Making Task (RDT)

After completing shaping, rats began training in the Risky Decision-Making Task (RDT). The beginning of each trial was marked by illumination of the house and trough lights, after which a poke into the lit trough caused either a single lever (free choice trials at the beginning of each new block) or both levers (forced choice trials) to extend. These levers were designated as “risky” or “safe”; pressing the “risky” lever resulted in delivery of three pellets along with risk of punishment (one sec foot shock) that increased with each subsequent block (0, 25, 50, 75, 100%). Selection of the “safe” lever resulted in one food pellet with no risk of shock. Choice of either lever caused immediate retraction of both levers, and after pellet collection all lights were extinguished and the trial proceeded to a 10s +/-4s inter trial interval (Gabriel et al., 2018; Figure 1a).

**Figure 1:**
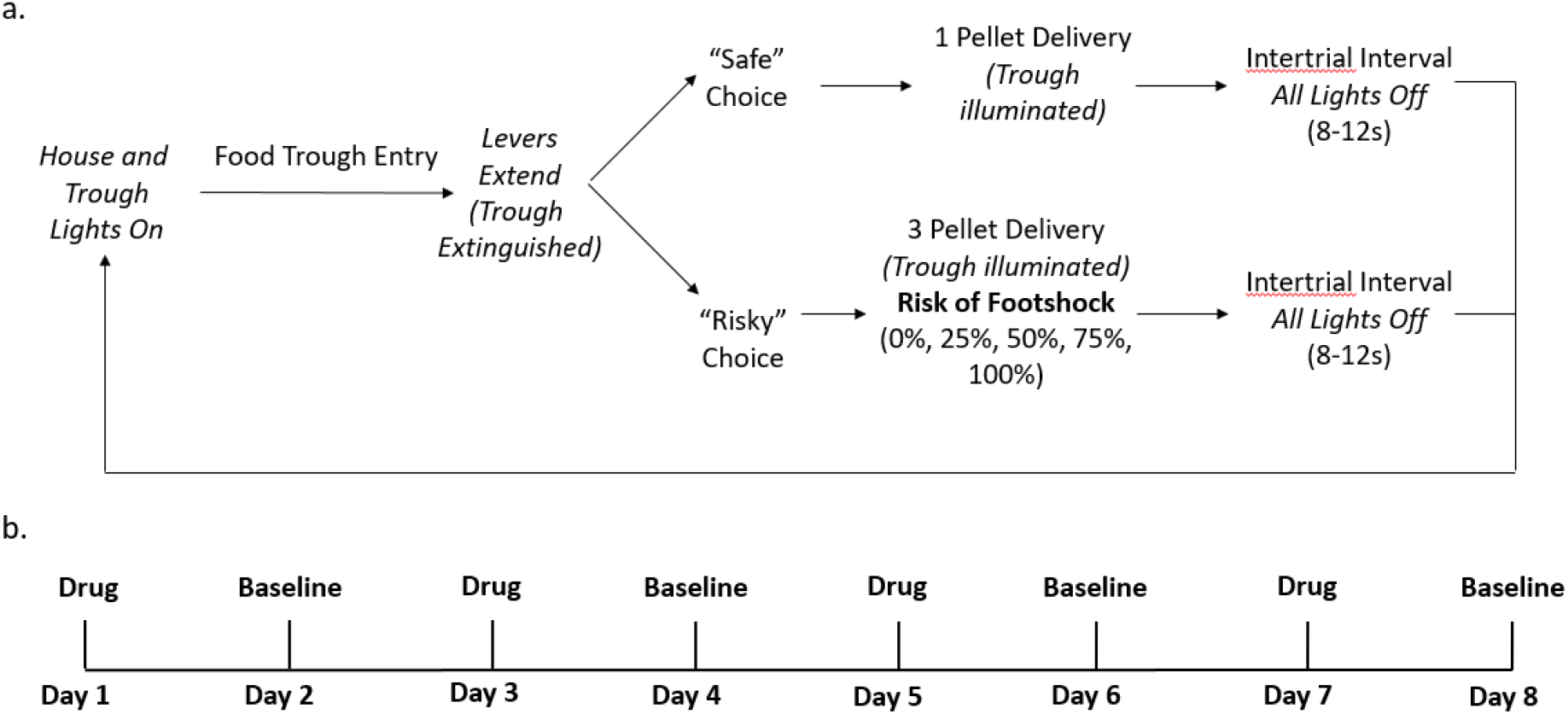
a) Risky Decision-making Task (RDT). This schematic depicts a single free choice trial of the RDT. B) Drug treatment procedure. For experiments one and two, subjects were given alternating drug treatment and drug-free baseline sessions for eight consecutive days. Dose order was counterbalanced for each drug. For experiment three, the treatment schedule followed a comparable pattern for four days (two drug doses/two baseline sessions).

Each session was divided into five 18 trial blocks for a total of 90 trials. Each block began with eight forced choice trials where each lever extended in a pseudorandom order, succeeded by ten free choice trials where both levers presented simultaneously (90 total trials per session). All parameters were equal across all blocks, except the risk of shock associated with the “risky” lever increased with each block.

Shock intensity was titrated for each individual subject to prevent floor or ceiling effects (Orsini and Simon, 2020). On the first session of RDT, shock amplitude was 0 mA (no shock), then was gradually increased on subsequent sessions by .02, .03, or .05 mA. If rats showed preference for 75% of the large-reward trials, shock was increased for the following session; if they preferred the large reward <25%, shock was decreased. Once discounting curves were stable for a minimum of two sessions, rats began drug injections.

### Drug Administration

The acute systemic effects of EGCG, caffeine, and combined EGCG and caffeine combination were examined across an eight-day schedule. For each drug regimen, the order of doses was counterbalanced across subjects to control for order or additive effects. All drugs were dissolved in .9% saline and administered at volume of 1 ml/kg via intraperitoneal injection.

For the EGCG regimen: subjects were administered EGCG (1.0, 2.0, 5.0 mg/kg) or saline vehicle on counterbalanced order on odd days (1,3,5,7), and non-drug baseline sessions on even days (2,4,6,8; Figure 1b). After each injection, rats remained in their home cage for 60 minutes, an absorption period sufficient to produce behavioral effects in rodents (Vignes et al., 2006). Caffeine (saline, 10.0, 17.0, 30.0 mg/kg) was administered on a similar eight-day schedule, using a 10-minute post injection absorption period prior to RDT (Diller et al., 2008). For the joint regimen, subjects were given an EGCG (5 mg/kg) and caffeine (2.5 mg/kg) combination that reflects the dose and ratio in a typical serving of green tea (Diller et al., 2008; Wu et al., 2012; Younes et al., 2018). For this four-day schedule, rats were given EGCG (5.0 mg/kg) 60 minutes before behavior and caffeine (2.5 mg/kg) 10 minutes before behavior, or two saline injections at comparable time intervals. Overall order of drug presentation was counterbalanced across rats, with a one-week washout period between each injection regimen.

### Data Analysis

Behavioral data was compiled using custom-made MATLAB scripts, and IBM SPSS Statistics 24 was utilized to conduct all statistical analyses. If Mauchly’s test of sphericity was violated, statistical reporting was adjusted using Greenhouse-Geisser values with degrees of freedom adjusted correspondingly. Rats that failed to select large reward ≤ 50% in block one (3 vs 1 pellets with no risk of punishment) during the saline session was removed from analysis.

After task acquisition, stable decision-making was assessed using a day x block repeated measures ANOVA, defined as 1.) a lack of effect of day, and 2.) a significant effect of block. Drug effects were analyzed using a drug x block repeated measures ANOVA. Separate ANOVAs were run to analyze the effects of each drug on behavioral performance during RDT. Furthermore, locomotion was recorded as means of observing effects of drugs on ambulation during safe and punished trials. To quantify this, time spent moving was averaged for both risky and safe trial types and compared using a one-way repeated measures ANOVA for each experiment.

Due to interruption of experiments related to laboratory shut down from the Covid-19 pandemic, a small subset of rats was given drug exposure that lasted longer than the standard 8-day protocol. In these instances, rats were tested to determine if their baseline was comparable to pre-pandemic conditions; if there was no significant difference in baseline, these subjects resumed drug exposure. Any subjects with a shift in baseline behavior were removed from the experiment. Baseline days following each drug administration were analyzed via dose x block ANOVA to determine that baseline behavior was stable.

## Results

### Experiment 1: Effects of EGCG on risky decision-making

Rats were tested during exposure to one of three doses of EGCG (1,2, or 5mg/kg) or saline vehicle. One rat was removed from analyses due to < 50% selection of the large reward in block one (n = 9). As expected, there was a main effect of risk block (*F*_(4, 32)_ = 19.428, *p* < .001) such that selection of the risky lever decreased with increasing risk across all doses (Figure 2a). While there was no main effect of EGCG dose on risk-taking (*F*_(3, 24)_ = 1.731, *p* = .187), there was a significant dose x block interaction (*F*_(12, 96)_ = 2.021, *p* = 030), such that the 5.0 mg/kg dose (but not the lower doses) reduced choice of the large, risky reward. We confirmed this by comparing only the 5.0mg/kg dose of EGCG with saline, which also yielded a significant treatment x block interaction (*F*_(4, 36)_ = 3.271, p = .022) such that EGCG did not alter decision-making with zero or low risk, but decreased risky reward choice as probability of punishment increased (Figure 2a). There were no differences between treatments when comparing either the 1 or 2 mg/kg doses with saline (*p* values >.2). Finally, there were no differences in reward choice between the four baseline days following drug administration (*F*_(1.612, 14.509)_ = .177, *p* = .794), suggesting that baseline risk preference remained stable throughout this experiment.

**Figure 2:**
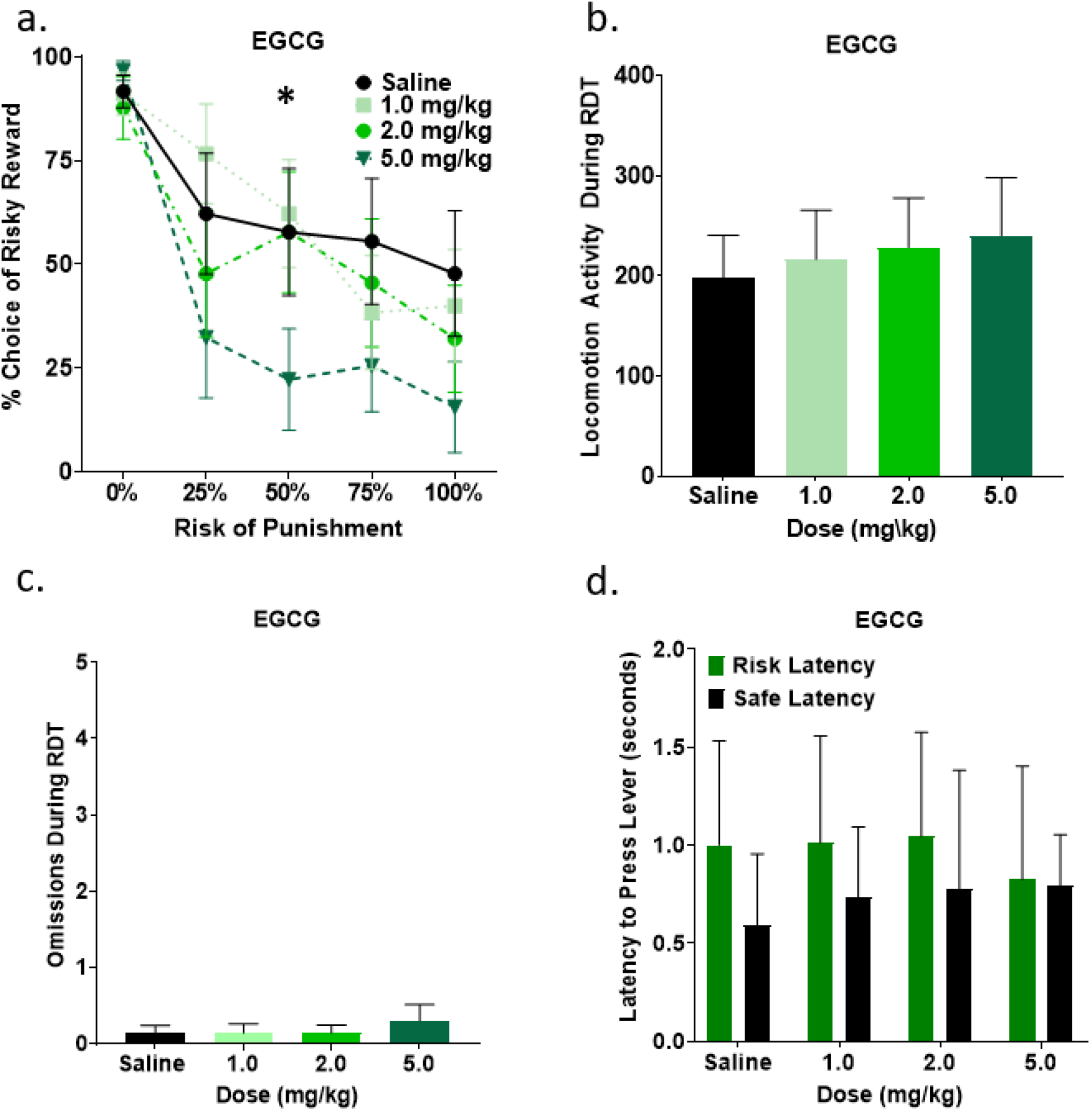
a) High dose EGCG reduced choice of the large, risky option. b) EGCG did not affect locomotor activity during the task. c) EGCG did not affect trial omissions. d). There was no difference in latency to choose the risky or safe reward across all doses. All panels depict mean ±SEM.

Next, we also assessed the effects of EGCG on other task-related behavioral measures. A one-way repeated measures ANOVA revealed that EGCG did not affect locomotion (*F*_(3, 27)_ = 1.231, *p* = .318; Figure 2b) or the number of trials omitted (*F*_(1.612, 14.512)_ = .338, *p* = .673; Figure 2c). Additionally, there was no effect of drug on latency to choose either the risky lever (*F*_(3, 27)_ = .449, *p =* .720) or safe lever (*F*_(3, 24)_ = 1.042, *p =* .392), indicating that exposure to EGCG did not influence rate of decision-making (Figure 2d). There was also no difference in latency to choose risky or safe choices (*F*_(1, 8)_ = 1.756, *p =* .222), or a choice type x EGCG dose interaction (*F*_(3, 24)_ = .165, *p =* .449). Collectively, these data suggest that the dose-dependent effects of EGCG were selective to risk-taking without affecting general measures of task engagement. Upon further analysis, there was a near significant difference in latency to pick the safe vs risky option after saline exposure (*t* (9) = 2.111, *p* = .064), but not after exposure to any dose of EGCG (*p*s > .05).

### Experiment 2: The Effects of Caffeine on Risky Decision-making

For this experiment, we investigated the acute effects of multiple doses of caffeine (10, 17, 30 mg/kg) or saline vehicle on risk-taking in RDT. There was a main effect of risk block (*F*_(4, 36)_ = 28.227, *p* < .001), such that choice of the large reward decreased as risk of punishment increased. However, there was no main effect of caffeine dose on risky decision-making (*F*_(1.443, 12.988)_ = 1.321, *p* = .289), nor was there a significant dose by block interaction (*F*_(5.090, 45.807)_ = .694, *p* = .633; Figure 3a). Upon visual inspection of the data, the low dose (10mg/kg) caused a qualitative increase in risk-taking; however, comparison between this dose and saline revealed no effect of drug (*F*_(1, 9)_ = 1.225, *p* = .297) and no drug x block interaction (*F*_(4, 36)_ = .448, *p* = .773). Thus, acute caffeine did not influence risky decision-making. There was no difference in risky decision-making between each of the four baseline days following drug administration (*F*_(3, 24)_ = .429, *p* = .734), indicating that risk preference was stable throughout the non-treatment portions of the injection regimen.

**Figure 3:**
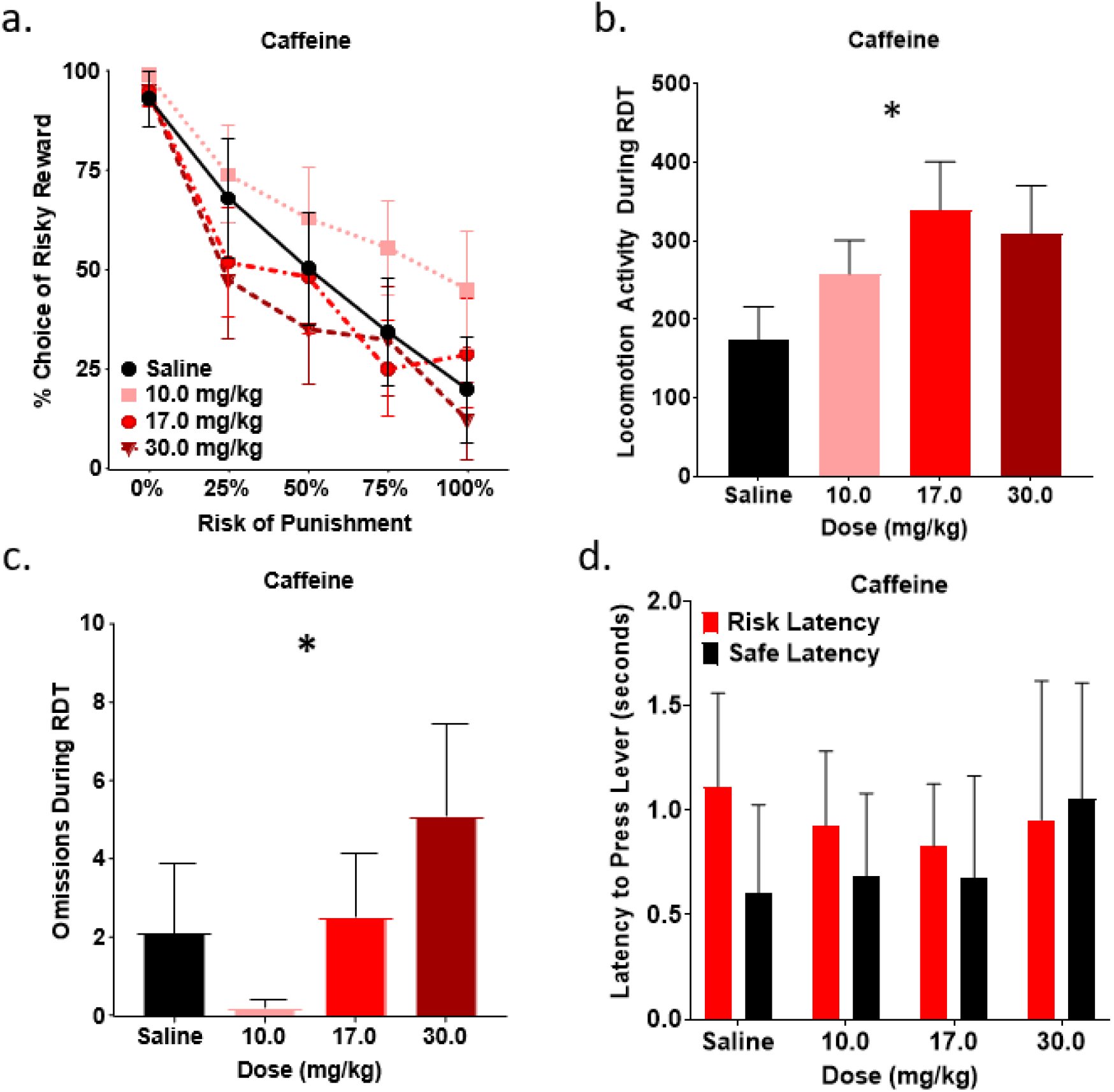
a) Caffeine did not alter risky decision-making. b) Caffeine caused a dose-dependent increase in locomotor activity during RDT. c) Caffeine exerted a dose-dependent effect on trial omissions, reducing omissions at low dose and increasing omissions at the highest dose. d) Latency to choose the safe option was shorter than the risky option. This was only significant after saline exposure. All panels depict mean ±SEM.

Next, we tested the effects of caffeine on locomotor activity. There was a significant effect of caffeine dose on locomotion (*F*_(3, 21)_ = 4.460, *p* = .014; Figure 3b), such that locomotion increased after caffeine exposure. Paired samples *t*-tests comparing individual doses of caffeine with saline revealed no significant increase in locomotion caused by the low dose (10mg/kg; *t* (9) = -1.208, *p* = .258), but significant increases in locomotion caused by the mid-range dose (17 mg/kg; *t* (9) = -2.537, *p* = .032) and high doses (30 mg/kg; *t* (9) = -3.204, *p* = .011). Therefore, caffeine caused a dose dependent elevation in locomotor activity.

A one-way repeated measures ANOVA revealed that caffeine dose had a significant effect on omissions (*F*_(3, 27)_ = 3.281, *p* = .036; Figure 3c). To further investigate this effect, each individual caffeine dose was compared to saline. The low and mid doses had no effect on omissions (low: *t* (9) = 1.046, *p* = .323; mid: *t* (9) = -.513, *p* = .620), but the high dose caused a significant increase in omissions (*t* (9) = -2.940, *p* = .016), suggesting that high dose caffeine reduces overall task engagement.

A mixed lever choice (risky vs safe) x caffeine dose ANOVA revealed a significant difference in latency to choose the risky vs safe lever (*F*_(1, 7)_ = 7.656, *p* = .028), such that rats selected the safe lever more rapidly than the risky lever (Figure 3d). Overall, caffeine did not affect latency to select either the risky (*F*_(3, 27)_ = .716, *p =* .551) or safe (*F*_(1.463, 10.239)_ = 1.237, *p =* .315) lever (Figure 3d). Upon further evaluation, rats chose the safe option more rapidly than the risky option when exposed to saline (*t* (8) = 3.052, *p* = .016), but not with any doses of caffeine (*p*s > .05).

### Experiment 3: The Effects of Co-administered EGCG and Caffeine

Finally, rats were administered both EGCG and caffeine to assess effects of exposure to the combined psychoactive ingredients in green tea. Two rats were excluded from analyses due to inability to achieve a stable baseline after laboratory shut down during the Covid-19 pandemic. There was a main effect of risk block (*F*_(2.283, 15.980)_ = 10.303, *p* = .001) showing that choice of the large reward decreased as risk of punishment increased. Despite this dose of EGCG reducing risk-taking in Experiment 1, combined EGCG and caffeine had no impact on risk-taking (*F* _(1, 7)_ = .366, *p* = .564; Figure 4a), and there was no drug x block interaction (*F*_(2.343, 16.398)_ = .673, *p =* .546). There was no difference in risk-taking between baseline days following drug administration, confirming that baseline risk-taking was consistent throughout the drug regimen (*F*_(1, 7)_ = .086, *p* = .778).

**Figure 4:**
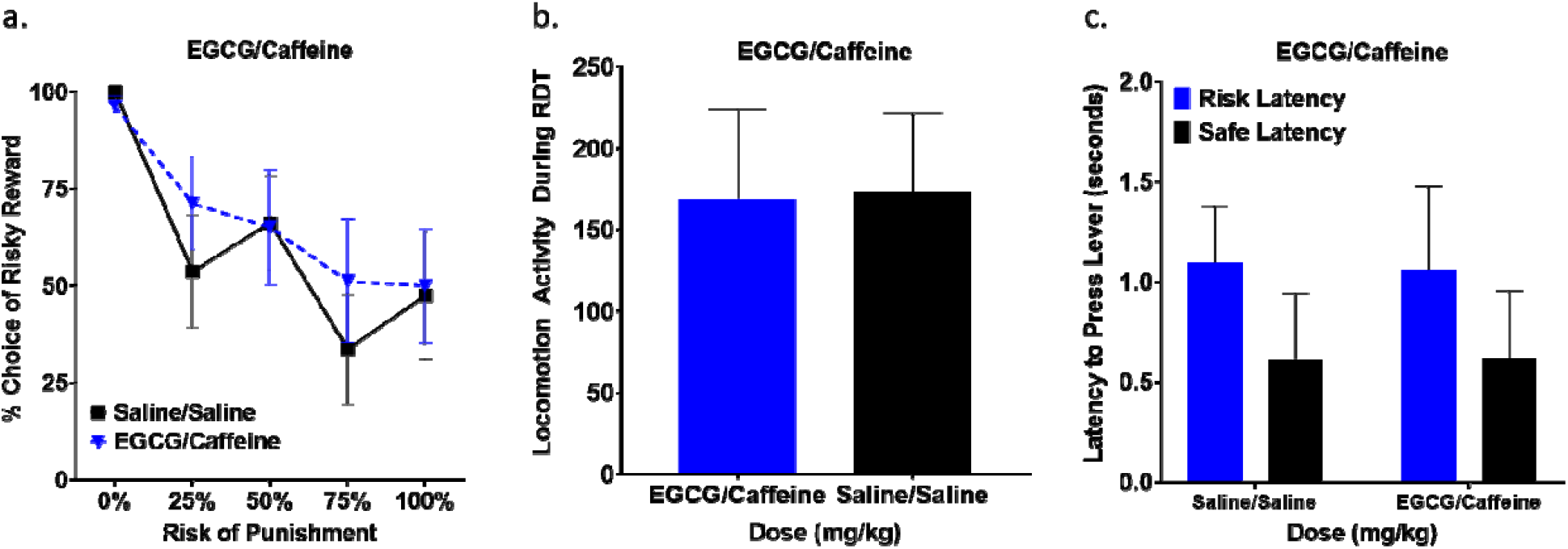
a) 5.0 mg/kg EGCG and 2.5mg/kg caffeine administered together had no effect on risk-taking. b) Locomotion was not influenced by combination EGCG and caffeine. c) Rats choose the safe reward more quickly than the risky option across both EGCG/caffeine and saline treatments. All panels depict mean ±SEM.

EGCG/caffeine had no effect on locomotion relative to saline (*F*_(1, 7)_ = .054, *p* = .823; Figure 4b), and there were no omissions following either injection schedule. A lever x dose repeated measures ANOVA revealed a significant difference in latency to choose the risky vs safe option (*F*_(1, 5)_ = 13.931, *p* = .014), such that rats chose the safe lever more quickly than the risky lever. There was no effect of drug on latency to make a choice (*F*_(1, 5)_ = .005, *p* = .946), nor was there a lever choice x drug interaction (*F*_(1, 5)_ = 4.013, *p* = .102; Figure 4c).

## Discussion

We observed that EGCG dose-dependently reduced risky decision-making without affecting other behavioral measures. Caffeine did not affect risk-taking, but increased locomotor activity and dose-dependently reduced completed trials. Interestingly, combining caffeine with EGCG in a ratio comparable to green tea consumption nullifies EGCG’s risk attenuating effects without affecting locomotion or other behavioral measures.

### Experiment 1: Effects of EGCG on risky decision-making

EGCG reduced risk-taking, but this effect required a relatively high dose of 5 mg/kg. This is comparable to 2-3 cups of green tea, with 3 cups being the amount consumed daily by the average tea drinker (Hu et al., 2018). As previously stated, EGCG induces anxiolytic and amnesic effects comparable to benzodiazepines (McCracken et al, 1990; Vignes et al., 2006). The risk aversion induced by EGCG is somewhat surprising, as the benzodiazepine diazepam caused the opposite effect of increased risk-taking (Mitchell et al., 2011). Diazepam and EGCG both modulate GABA_A_ receptor activity at the benzodiazepine site (Campbell et al., 2004; Hanrahan et al., 2011; Calcaterra and Barrow, 2014); diazepam is well-established as an allosteric modulator that increases GABA-induced chloride influx (Calcaterra and Barrow, 2014), whereas the pharmacodynamics of EGCG at GABA receptors are less well-defined (Hanrahan et al., 2011). Based on the diverging effects of EGCG and diazepam on risk-taking, it is likely that 1.) there are critical yet unclear differences in pharmacodynamics between these drugs that evoke different effects on decision-making, and 2.) risk-taking is not mediated unidirectionally by anxiety, as these drugs both exert anxiolytic effects, yet opposing effects on risk-taking.

Green tea polyphenols such as EGCG increase synaptic dopamine by reducing dopamine reuptake (Pan et al., 2003; Li et al., 2006). Previous research has shown that amphetamine, which also inhibits dopamine reuptake (Faraone, 2018), and agonists at D2 receptors reduce risk taking in a manner similar to EGCG (Setlow et al., 2009; Simon et al., 2011; Orsini et al., 2016, 2019). Thus, it is possible that EGCG induces risk-aversion through increased dopamine receptor activation.

Due to the use of footshock in the RDT, it is possible that EGCG’s effects on risk are related to analgesia, as green tea polyphenols can cause antinociception (Lee et al., 2018). However, EGCG reduced choice of the punished reward, whereas analgesia would be expected to increase tolerance of the footshock. Moreover, morphine had no effect on risk-taking in RDT (Mitchell et al., 2011), and RDT performance is uncorrelated with shock sensitivity (Simon et al., 2011), suggesting that risk-taking is not strongly modulated by pain tolerance. Additionally, EGCG did affect overall locomotion, trial omissions, or latencies to make either risky or safe decisions. This indicates that EGCG’s effects on risk-taking were likely not related to ambulation, reduced task engagement, or cognitive disruption related to immediate reward preference. It is also unlikely that EGCG caused reduced choice in the large reward due to a gross reduction in motivation, as choice of the large reward in block one (0% shock) was unaffected.

### Experiment 2: The Effects of Caffeine on Risky Decision-making

Caffeine has been shown to affect multiple forms of economic decision-making (Smit and Rogers, 2000; Diller et al., 2008; Temple et al., 2017). However, acute caffeine had no effects on risky decision-making. This contrasts with the risk-aversion observed after exposure to the psychostimulant drugs amphetamine and nicotine, suggesting that risk aversion does not generalize across all psychostimulants. Caffeine does not typically affect risk-taking in humans (Killgore et al., 2007), but can restore risk-taking to baseline levels after long-term sleep deprivation (Killgore et al., 2011). All experiments here were performed during the dark cycle to enhance wakefulness; it is possible that repeating the current experiments during the light cycle or after sleep restriction would enhance caffeine’s effects on risk-taking. Notably, 12 hours of sleep deprivation on rats does not affect probabilistic discounting (Leenaars et al., 2021); thus, longer sleep restriction may be necessary to affect baseline risk-taking sufficiently to enable assessment of the restorative effects of caffeine or other psychostimulants.

Caffeine is a known psychostimulant that causes increased locomotor activity similar to that of other stimulants such as cocaine and amphetamine via dopamine release (Garrett and Griffiths, 1997). Caffeine’s stimulating effects on locomotion in this study are consistent with previous research (Cauli and Morelli, 2005); this hyperactivity demonstrates that despite caffeine’s lack of effects on decision-making, the doses used here were sufficient to affect behavior. Furthermore, a dose dependent increase in trial omissions was also observed with the high dose of caffeine (30mg/kg), indicating that high dose caffeine reduces task engagement in similar fashion to dopaminergic psychostimulants (Simon et al., 2009). Interestingly, a comparable dose did not increase omissions in a delay discounting task (Diller et al., 2008), suggesting that caffeine only reduces task engagement during decision-making with aversive outcomes.

We observed a difference in latency to select the safe vs risky reward, such that safe decisions were faster than risky decisions. This is consistent with previous reports of RDT, suggesting that conditioned fear evoked by the punishment-associated risky lever may cause either brief freezing behavior or longer deliberation prior to choice (Orsini et al., 2015). Notably, this difference was only significant after saline treatment, suggesting that caffeine may have reduced the latency to choose the risky reward despite not impacting risk preference. Notably, while this difference between risky and safe decision-making latency was also observed during the caffeine/EGCG combination experiment, it was not observed during the EGCG experiment, although choice of the risky reward was qualitatively slower than the safe reward after saline exposure.

### Experiment 3: The Effects of Co-administered Caffeine and EGCG

Administered individually, EGCG dose-dependently reduces risk-taking, and caffeine does not affect risk-taking. Interestingly, when EGCG and caffeine were both administered, risk-taking was unaffected. Rather than use the higher doses of caffeine from experiment two, we selected specific doses for this experiment to approximate daily consumption of EGCG and caffeine by the average green tea consumer (Younes et al., 2018). Despite this relatively low dose, caffeine exposure was sufficient to block the effects of EGCG on risk-taking.

Joint administration of a psychostimulant and sedative can cause opposing effects on arousal, which can mask the individual stimulant/sedative effects of each drug (Leri et al., 2003; McKetin et al., 2015). Han et al. (2011, 2016) found that EGCG is protective against increases in blood pressure, heart rate, as well as blood levels of adrenaline, noradrenaline, and dopamine induced by caffeine intake. However, little is known about the combined effects of stimulant and sedative drugs on risky decision-making. In humans, results have been variable: consumers of combined alcohol and energy drinks typically report greater risk-taking than alcohol consumers, but repeated measures studies have revealed that energy drink consumption can reduced alcohol-induced risk-taking (Peacock et al., 2014). The results here are more aligned with the latter report, with caffeine eliminating the effects of EGCG (which exerts mild sedative and hypnotic effects (Adachi et al., 2006) on risk-taking, although alcohol and EGCG exerted opposing effects on risk-taking.

Notably, EGCG and caffeine were not administered simultaneously, but were given separately at different time intervals, with EGCG given one hour before testing and caffeine 10 minutes prior to testing RDT. These intervals were selected to enable enough absorption of each drug to affect behavior in rodents (Diller et al., 2008; Wu et al., 2012; Younes et al., 2018). Furthermore, despite the lengthy absorption period, this interval was sufficient to produce effects of EGCG on risk-taking in Experiment 1 that persisted the entire session. When drinking green tea, EGCG and caffeine would be consumed simultaneously, so it is possible that this difference in absorption may affect the joint influence of these compounds on cognition and arousal.

## Conclusion

Despite the worldwide prevalence of green tea consumption and the evidence of therapeutic effects of EGCG, there is a lack of research on the effects of EGCG on cognition and behavior. This experiment is the first to examine the effects of EGCG on risk-taking, as well as the acute effects of caffeine and joint EGCG/caffeine. The results suggest EGCG as a potential therapeutic treatment for psychological disorders that induce compulsive risky decision-making, such as bipolar disorder, schizophrenia, and substance use disorder (Reddy et al., 2013; Chen et al., 2020). Notably, these effects are nullified when EGCG is given in conjunction with caffeine, suggesting that EGCG must be isolated from green tea to impact cost-benefit decision-making. More research on the effects of EGCG on decision-making relevant brain circuitry is necessary to uncover why EGCG influences risk-taking, and why its effects contrast with other anxiolytic drugs such as diazepam and alcohol (Mitchell et al., 2011).

